# Four novel *Caudoviricetes* bacteriophages isolated from Baltic Sea water infect colonizers of *Aurelia aurita*

**DOI:** 10.1101/2023.06.01.543199

**Authors:** M. Stante, N. Weiland-Bräuer, U. Repnik, A. Werner, C. M. Chibani, M. Bramkamp, R. A. Schmitz

**Affiliations:** Institute for General Microbiology, Christian Albrechts University, Kiel, Germany; Central Microscopy Facility, Christian-Albrechts-University Kiel, Kiel, Germany

**Keywords:** bacteriophage, lytic cycle, Baltic Sea, *Caudoviricetes*, *Pseudomonas*, *Citrobacter*

## Abstract

The moon jellyfish *Aurelia aurita* is associated with a highly diverse microbiota changing with provenance, tissue, and life stage. While the crucial relevance of bacteria to host fitness is well known, bacteriophages have often been neglected. Here, we aimed to isolate lytic phages targeting bacteria that are part of the *A. aurita*-associated microbiota. Four phages (*Pseudomonas* phage BSwM KMM1, *Citrobacter* phages BSwM KMM2- BSwM KMM4) were isolated from the Baltic Sea water column and characterized. Phages KMM2/3/4 infected representatives of *Citrobacter*, *Shigella, and Escherichia* (*Enterobacteriaceae),* whereas KMM1 showed a remarkably broad host range, infecting Gram-negative *Pseudomonas* as well as Gram-positive *Staphylococcus*. All phages showed short latent periods (around 30 min), large burst sizes (mean of 128 pfu/mL), and high efficiency of plating (EOP > 0.5), demonstrating decent virulence, efficiency, and infectivity. Transmission electron microscopy and viral genome analysis revealed that all phages are novel species and belong to the tailed, linear double-stranded DNA phage families *Siphovirus* (KMM3) and *Myovirus* (KMM1/2/4), with genome sizes between 50 - 138 kbp. Those isolates now allow manipulation of the *A. aurita*-associated microbiota and will provide new insights into phage impact on the multicellular host.

## Introduction

A metaorganism is defined as the collective entity formed by a multicellular host and the diverse community of microorganisms that live in and on it (Bosch & McFall-Ngai, 2011). The metaorganism concept recognizes that hosts and their associated microbiota, including bacteria, archaea, fungi, and viruses, are interdependent and function as a solitary biological system, with the combined genetic repertoire of the metaorganism playing a key role in its overall health and survival (Ainsworth et al., 2020; Jaspers et al., 2019). The host organism provides habitat and nutrients for the microorganisms, while the microorganisms perform various functions that benefit the host, such as aiding digestion (Ley et al., 2008), modulating the immune system (Esser et al., 2019), and protecting against pathogens (He et al., 2020). Metaorganism research aims to understand the complex interactions between host organisms and their associated microorganisms and how these interactions impact the health and functioning of the metaorganism (Bang et al., 2018). Numerous studies on the diversity and impact of bacteria on a host have been published in recent years (Masuzzo et al., 2020; Wong et al., 2016; Xiao et al., 2022). However, other key players, such as bacteriophages, have often been neglected (Weiland-Bräuer et al., 2015; Weiland-Bräuer et al., 2020; Weiland-Bräuer et al., 2020). Bacteriophages, or phages for short, are viruses that infect bacteria. They are widespread in nature and are considered Earth’s most abundant biological entities (Batinovic et al., 2019; Salmond & Fineran, 2015). Phages can be classified based on their replication cycle, morphology, nucleic acid type, and host range. Particularly, their replication mode, such as lytic or lysogenic, is an essential characteristic (Hobbs & Abedon, 2016). Lytic phages cause lysis of the host bacterium (Węgrzyn, 2022), whereas lysogenic phages integrate their genetic material into the host bacterium’s genome and can remain dormant until conditions favor their activation into the lytic cycle (Bertani, 1951; Hobbs & Abedon, 2016; Lwoff, 1953). In recent years, the interest in the role of phages in host-associated microbiomes has increased (Adair & Douglas, 2017; Greenspan et al., 2020; Parfrey et al., 2018; Welsh et al., 2016). The presence and activity of lysogenic and lytic phages in microbiomes can significantly impact the microbial community’s composition and function and consequently affect host health (Mirzaei & Maurice, 2017; Zuppi et al., 2022). The lysogenic phage (prophage) replicates with the host genome and is transmitted to daughter cells during cell division (Canchaya et al., 2003). Integrated into the host genome, they can impact the microbiome by altering the gene expression of the host bacterium, leading to changes in its physiology and metabolism (Howard-Varona et al., 2017). Additionally, the prophage can provide additional new genes to the host bacterium and confer advantages, such as antibiotic resistance or enhanced metabolic capabilities (so-called auxiliary genes), but diseases caused directly by prophage-encoded virulence factors, such as botulism, diphtheria, and cholera, are also known to occur (Abedon & LeJeune, 2005; Pfeifer et al., 2022; Thompson et al., 2011; Tuttle & Buchan, 2020). Lytic phages have the ability to reduce the population of the host bacteria in the microbiome (Miller-Ensminger et al., 2018), which in turn can have a significant impact on the microbiome’s structure and function, as it changes the relative abundance of different bacterial species and alter the metabolic activities of the community. Well known examples include phages that have been shown to play an important role in the microbiomes of many invertebrates (Kirsch et al., 2021; Zhou et al., 2022), including sponges (Schmittmann et al., 2020), and within the squid’s light organ (Lynch et al., 2022).

One approach to studying the role of phages in metaorganisms is metagenomic sequencing to analyze the phage populations in the microbiome (Benler et al., 2021; Casas & Rohwer, 2007; Deines et al., 2017). Such studies can provide insights into the diversity, abundance, infection cycles (lytic or lysogenic), and activity of the phages and their interactions with the bacterial populations in the microbiome (Federici et al., 2021; Mirzaei & Maurice, 2017). However, functional studies on the role of phages in microbiomes require the isolation of phages, which are, due to methodological challenges, primarily focusing on lytic phages (Hyman & Abedon, 2009; Salmond & Fineran, 2015). Lytic phages can be isolated from various sources, including soil, water, and sewage, using a cultivation-dependent enrichment and isolation procedure. The cultivation-dependent approach includes sample collection and preparation, enrichment using selected host bacteria, isolation, and propagation (Jofre & Muniesa, 2020). Isolated novel phages can subsequently be characterized to provide a comprehensive understanding of the phage’s morphology, genetics, and behavior, which can be useful for functional studies to elucidate their role in the metaorganism, and applications in phage therapy and biotechnology (Hyman, 2019).

Although phages have been recognized as important players in microbial communities, the research on the role of phages in metaorganisms is still in its infancy. Consequently, this study aimed to isolate lytic phages from Baltic Sea water infecting bacterial colonizers of our metaorganism model, the moon jellyfish *Aurelia aurita. A. aurita* is a Cnidarian jellyfish found in many parts of the world’s oceans, particularly common in coastal areas and estuaries (Hubot et al., 2021). *A. aurita* has a relatively short lifespan as a mature medusa, usually living only for a few months to a year (Ceh et al., 2015). During this time, *A. aurita* undergoes a complex life cycle that includes both asexual and sexual reproduction (Ceh et al., 2015). *A. aurita* is associated with a highly diverse microbiota depending on its provenance, tissue, and life stage (Weiland-Bräuer et al., 2015). This specific microbiota is crucial for the survival, growth, and asexual reproduction of the host (Weiland-Bräuer et al., 2020). A total of 132 bacterial representatives of the associated microbiota were derived from different sub-populations and life stages of *A. aurita* (Weiland-Bräuer et al., 2020), representing different taxonomic groups including Proteobacteria (*Pseudomonas*, *Alteromonas*, *Pseudoalteromonas*, *Vibrio*, *Paracoccus*, *Ruegeria*, *Shewanella*, *Sulfitobacter*), Actinomycetota (*Rhodococcus*, *Brevibacterium*, *Microbacterium*, *Micrococcus*), Bacilli (*Bacillus*, *Enterococcus*, *Streptococcus*), and Flavobacteriia (*Chryseobacterium*, *Maribacter*, *Olleya*). Though the importance and impact of bacteria on the health and fitness of *A. aurita* has already been demonstrated (Jensen et al., 2023; Weiland-Bräuer et al., 2015; Weiland-Bräuer et al., 2020; Weiland-Bräuer et al., 2020), bacteriophages have not previously been considered in this context. The present study reports on the morphological, microbiological, and genomic characterization of four newly isolated lytic phages, (*Pseudomonas* phage BSwM KMM1, *Citrobacter* phages BSwM KMM2, BSwS KMM3, and BSwM KMM4) infecting colonizers of *A. auritás* microbiota.

## Material and Methods

### Bacterial strains and growth conditions

The bacterial strains used in this study are listed in **Table 1** with their respective culture media and growth conditions. Bacterial strains of marine origin were grown in Marine Bouillon (MB; 10 g/L yeast extract, 10 g/L peptone (Carl Roth, Karlsruhe, Germany), 30 practical salinity units (PSU) Tropical Marine Salts, pH 7.3). Other bacterial strains were cultivated, following the recommendations, in Trypticase Soy Yeast Broth (TSYB, Carl Roth, Karlsruhe, Germany), Caso-Bouillon (CASO, Carl Roth, Karlsruhe, Germany), Lysogeny Broth (LB, Carl Roth, Karlsruhe, Germany), Nutrient Broth (NB, Carl Roth, Karlsruhe, Germany) according to the DSMZ (German Collection of Microorganisms and Cell Cultures GmbH).

**Table 1.** Bacterial strains used in this study. Bacterial strains were isolated in the study (Weiland-Bräuer et al., 2020). Isolates are sorted by the last column and phylum level. The column [use in this study] refers to the use of the strains in the study. If not stated differently, the listed numbers in column [reference] reflect NCBI Accession Nos.

| strain no. | strain | reference | phylum | class | order | family | source | growth medium | growth temp. | use in this study |
| --- | --- | --- | --- | --- | --- | --- | --- | --- | --- | --- |
| 74 | *Micrococcus luteus* | MK967048.1 | Actinomycetota | Actinomycetia | Micrococcales | Micrococcaceae | *A. aurita* polyp Baltic Sea husbandry | Marine Bouillon | 30°C | enrichment/ first screening |
| 75 | *Arthrobacter sp.* | MK967049.1 | Actinomycetota | Actinomycetia | Micrococcales | Micrococcaceae | *A. aurita* polyp Baltic Sea husbandry | Marine Bouillon | 30°C | enrichment/ first screening |
| 83 | *Gordonia terrae* | MK967057.1 | Actinomycetota | Actinomycetia | Mycobacteriales | Gordoniaceae | *A. aurita* polyp Baltic Sea husbandry | Marine Bouillon | 30°C | enrichment/ first screening |
| 15 | *Sulfitobacter sp.* | MK967015.1 | Pseudomonadota | Alphaproteobacteria | Rhodobacterales | Rhodobacteraceae | *A. aurita* medusa Baltic Sea | Marine Bouillon | 30°C | enrichment/ first screening |
| 20 | *Sulfitobacter pontiacus* | MK967020.1 | Pseudomonadota | Alphaproteobacteria | Rhodobacterales | Rhodobacteraceae | *A. aurita* medusa Baltic Sea husbandry | Marine Bouillon | 30°C | enrichment/ first screening |
| 23 | *Sulfitobacter sp.* | MK967023.1 | Pseudomonadota | Alphaproteobacteria | Rhodobacterales | Rhodobacteraceae | *A. aurita* medusa Baltic Sea husbandry | Marine Bouillon | 30°C | enrichment/ first screening |
| 69 | *Rhodobacter sp.* | MK967043.1 | Pseudomonadota | Alphaproteobacteria | Rhodobacterales | Rhodobacteraceae | *M. leidy* Baltic Sea husbandry | Marine Bouillon | 30°C | enrichment/ first screening |
| 78 | *Sulfitobacter sp.* | MK967052.1 | Pseudomonadota | Alphaproteobacteria | Rhodobacterales | Rhodobacteraceae | *A. aurita* polyp Baltic Sea husbandry | Marine Bouillon | 30°C | enrichment/ first screening |
| 86 | *Ruegeria sp.* | MK967060.1 | Pseudomonadota | Alphaproteobacteria | Rhodobacterales | Rhodobacteraceae | *A. aurita* polyp Baltic Sea husbandry | Marine Bouillon | 30°C | enrichment/ first screening |
| 89 | *Ruegeria sp.* | MK967063.1 | Pseudomonadota | Alphaproteobacteria | Rhodobacterales | Rhodobacteraceae | *A. aurita* polyp Baltic Sea husbandry | Marine Bouillon | 30°C | enrichment/ first screening |
| 100 | *Sulfitobacter sp.* | MK967074.1 | Pseudomonadota | Alphaproteobacteria | Rhodobacterales | Rhodobacteraceae | *A. aurita* polyp North Sea husbandry | Marine Bouillon | 30°C | enrichment/ first screening |
| 117 | *Ruegeria mobilis* | MK967091.1 | Pseudomonadota | Alphaproteobacteria | Rhodobacterales | Rhodobacteraceae | *A. aurita* polyp North Atlantic husbandry | Marine Bouillon | 30°C | enrichment/ first screening |
| 147 | *Phaeobacter gallaeciensis* | MK967120.1 | Pseudomonadota | Alphaproteobacteria | Rhodobacterales | Rhodobacteraceae | Artificial Seawater 18 PSU | Marine Bouillon | 30°C | enrichment/ first screening |
| 188 | *Sulfitobacter pseudonitzschiae* | MK967160.1 | Pseudomonadota | Alphaproteobacteria | Rhodobacterales | Rhodobacteraceae | Artificial Seawater 30 PSU | Marine Bouillon | 30°C | enrichment/ first screening |
| 13 | *Bacillus cereus* | MK967013.1 | Bacillota | Bacilli | Bacillales | Bacillaceae | *A. aurita* medusa Baltic Sea | Marine Bouillon | 30°C | enrichment/ first screening |
| 16 | *Bacillus sp.* | MK967016.1 | Bacillota | Bacilli | Bacillales | Bacillaceae | *A. aurita* medusa Baltic Sea | Marine Bouillon | 30°C | enrichment/ first screening |
| 17 | *Bacillus cereus* | MK967017.1 | Bacillota | Bacilli | Bacillales | Bacillaceae | *A. aurita* medusa Baltic Sea | Marine Bouillon | 30°C | enrichment/ first screening |
| 19 | *Bacillus sp.* | MK967019.1 | Bacillota | Bacilli | Bacillales | Bacillaceae | *A. aurita* medusa Baltic Sea husbandry | Marine Bouillon | 30°C | enrichment/ first screening |
| 76 | *Bacillus weihenstephanensis* | MK967050.1 | Bacillota | Bacilli | Bacillales | Bacillaceae | *A. aurita* polyp Baltic Sea husbandry | Marine Bouillon | 30°C | enrichment/ first screening |
| 85 | *Staphylococcus warneri* | MK967059.1 | Bacillota | Bacilli | Bacillales | Staphylococcaceae | *A. aurita* polyp Baltic Sea husbandry | Marine Bouillon | 30°C | enrichment/ first screening |
| 88 | *Staphylococcus sp.* | MK967062.1 | Bacillota | Bacilli | Bacillales | Staphylococcaceae | *A. aurita* polyp Baltic Sea husbandry | Marine Bouillon | 30°C | enrichment/ first screening |
| 73 | *Enterococcus casseliflavus* | MK967047.1 | Bacillota | Bacilli | Lactobacillales | Enterococcaceae | *A. aurita* polyp Baltic Sea husbandry | Marine Bouillon | 30°C | enrichment/ first screening |
| 24 | *Maribacter* sp. | MK967024.1 | Bacteroidota | Flavobacteriia | Flavobacteriales | Flavobacteriaceae | *A. aurita* medusa Baltic Sea husbandry | Marine Bouillon | 30°C | enrichment/ first screening |
| 57 | *Olleya marilimosa* | MK967032.1 | Bacteroidota | Flavobacteriia | Flavobacteriales | Flavobacteriaceae | *M. leidy* Baltic Sea | Marine Bouillon | 30°C | enrichment/ first screening |
| 79 | *Olleya* sp. | MK967053.1 | Bacteroidota | Flavobacteriia | Flavobacteriales | Flavobacteriaceae | *A. aurita* polyp Baltic Sea husbandry | Marine Bouillon | 30°C | enrichment/ first screening |
| 181 | *Chryseobacterium* sp. | MK967154.1 | Bacteroidota | Flavobacteriia | Flavobacteriales | Weeksellaceae | Artificial Seawater 30 PSU | Marine Bouillon | 30°C | enrichment/ first screening |
| 257 | *Chryseobacterium* sp. | MK967218.1 | Bacteroidota | Flavobacteriia | Flavobacteriales | Weeksellaceae | *M. leidy* Baltic Sea husbandry | Marine Bouillon | 30°C | enrichment/ first screening |
| 22 | *Pseudolateromonas* sp. | MK967022.1 | Pseudomonadota | Gammaproteobacteria | Alteromonadales | Pseudoalteromonadaceae | *A. aurita* medusa Baltic Sea husbandry | Marine Bouillon | 30°C | enrichment/ first screening |
| 91 | *Pseudoalteromonas prydzensis* | MK967065.1 | Pseudomonadota | Gammaproteobacteria | Alteromonadales | Pseudoalteromonadaceae | *A. aurita* polyp Baltic Sea husbandry | Marine Bouillon | 30°C | enrichment/ first screening |
| 101 | *Pseudoalteromonas issachenkonii* | MK967075.1 | Pseudomonadota | Gammaproteobacteria | Alteromonadales | Pseudoalteromonadaceae | *A. aurita* polyp North Sea husbandry | Marine Bouillon | 30°C | enrichment/ first screening |
| 167 | *Pseudoalteromonas* sp. | MK967140.1 | Pseudomonadota | Gammaproteobacteria | Alteromonadales | Pseudoalteromonadaceae | Artificial Seawater 18 PSU | Marine Bouillon | 30°C | enrichment/ first screening |
| 203 | *Pseudoalteromonas espejiana* | MK967174.1 | Pseudomonadota | Gammaproteobacteria | Alteromonadales | Pseudoalteromonadaceae | Artificial Seawater 30 PSU | Marine Bouillon | 30°C | enrichment/ first screening |
| 219 | *Pseudoalteromonas tunicata* | MK967188.1 | Pseudomonadota | Gammaproteobacteria | Alteromonadales | Pseudoalteromonadaceae | *M. leidy* Baltic Sea | Marine Bouillon | 30°C | enrichment/ first screening |
| 224 | *Pseudoalteromonas lipolytica* | MK967191.1 | Pseudomonadota | Gammaproteobacteria | Alteromonadales | Pseudoalteromonadaceae | *M. leidy* Baltic Sea | Marine Bouillon | 30°C | enrichment/ first screening |
| 105 | *Shewanella basaltis* | MK967079.1 | Pseudomonadota | Gammaproteobacteria | Alteromonadales | Shewanellaceae | *A. aurita* polyp North Sea husbandry | Marine Bouillon | 30°C | enrichment/ first screening |
| 21 | *Cobetia amphilecti* | MK967021.1 | Pseudomonadota | Gammaproteobacteria | Oceanospirillales | Halomonadaceae | *A. aurita* medusa Baltic Sea husbandry | Marine Bouillon | 30°C | enrichment/ first screening |
| 55 | *Marinomonas hwangdonensis* | MK967030.1 | Pseudomonadota | Gammaproteobacteria | Oceanospirillales | Oceanospirillaceae | *M. leidy* Baltic Sea | Marine Bouillon | 30°C | enrichment/ first screening |
| 222 | *Marinomonas pontica* | MK967189.1 | Pseudomonadota | Gammaproteobacteria | Oceanospirillales | Oceanospirillaceae | *M. leidy* Baltic Sea | Marine Bouillon | 30°C | enrichment/ first screening |
| 262 | *Oceanospirillaceae bacterium* | MK967222.1 | Pseudomonadota | Gammaproteobacteria | Oceanospirillales | Oceanospirillaceae | *M. leidy* Baltic Sea | Marine Bouillon | 30°C | enrichment/ first screening |
| 11 | *Pseudomonas sp.* | MK967012.1 | Pseudomonadota | Gammaproteobacteria | Pseudomonadales | Pseudomonadaceae | *A. aurita* medusa Baltic Sea | Marine Bouillon | 30°C | enrichment/ first screening |
| 90 | *Pseudomonas putida* | MK967064.1 | Pseudomonadota | Gammaproteobacteria | Pseudomonadales | Pseudomonadaceae | *A. aurita* polyp Baltic Sea husbandry | Marine Bouillon | 30°C | enrichment/ first screening |
| 92 | *Pseudomonas putida* | MK967066.1 | Pseudomonadota | Gammaproteobacteria | Pseudomonadales | Pseudomonadaceae | *A. aurita* polyp Baltic Sea husbandry | Marine Bouillon | 30°C | enrichment/ first screening |
| 93 | *Pseudomonas sp.* | MK967067.1 | Pseudomonadota | Gammaproteobacteria | Pseudomonadales | Pseudomonadaceae | *A. aurita* polyp Baltic Sea husbandry | Marine Bouillon | 30°C | enrichment/ first screening |
| 94 | *Pseudomonas sp.* | MK967068.1 | Pseudomonadota | Gammaproteobacteria | Pseudomonadales | Pseudomonadaceae | *A. aurita* polyp Baltic Sea husbandry | Marine Bouillon | 30°C | enrichment/ first screening |
| 132 | *Pseudomonas fluorescens* | MK967106.1 | Pseudomonadota | Gammaproteobacteria | Pseudomonadales | Pseudomonadaceae | Artificial Seawater 18 PSU | Marine Bouillon | 30°C | enrichment/ first screening |
| 196 | *Pseudomonas syringae* | MK967168.1 | Pseudomonadota | Gammaproteobacteria | Pseudomonadales | Pseudomonadaceae | Artificial Seawater 30 PSU | Marine Bouillon | 30°C | enrichment/ first screening |
| 77 | *Vibrio anguillarum* | MK967051.1 | Pseudomonadota | Gammaproteobacteria | Vibrionales | Vibrionaceae | *A. aurita* polyp Baltic Sea husbandry | Marine Bouillon | 30°C | enrichment/ first screening |
| 80 | *Vibrio anguillarum* | MK967054.1 | Pseudomonadota | Gammaproteobacteria | Vibrionales | Vibrionaceae | *A. aurita* polyp Baltic Sea husbandry | Marine Bouillon | 30°C | enrichment/ first screening |
| 18 | *Staphylococcus aureus* | OQ398157 | Bacillota | Bacilli | Bacillales | Staphylococcaceae | *A. aurita* medusa Baltic Sea husbandry | Marine Bouillon | 30°C | enrichment/ first screening |
| 134 | *Staphylococcus aureus* | OQ398164 | Bacillota | Bacilli | Bacillales | Staphylococcaceae | Artificial Seawater 18 PSU | Marine Bouillon | 30°C | enrichment/ first screening |
| 87 | *Staphylococcus aureus* | OQ398160 | Bacillota | Bacilli | Bacillales | Staphylococcaceae | *A. aurita* polyp Baltic Sea husbandry | Marine Bouillon | 30°C | enrichment/ first screening |
| 14 | *Staphylococcus warneri* | OQ398156 | Bacillota | Bacilli | Bacillales | Staphylococcaceae | *A. aurita* medusa Baltic Sea | Marine Bouillon | 30°C | enrichment/ first screening |
| 6 | *Citrobacter freundii* | OQ398153 | Pseudomonadota | Gammaproteobacteria | Enterobacterales | Enterobacteriaceae | *A. aurita* medusa Baltic Sea | Marine Bouillon | 30°C | enrichment/ first screening/ host range |
| 7 | *Citrobacter sp.* | OQ398154 | Pseudomonadota | Gammaproteobacteria | Enterobacterales | Enterobacteriaceae | *A. aurita* medusa Baltic Sea | Marine Bouillon | 30°C | enrichment/ first screening/ host range |
| 8 | *Pseudomonas sp.* | MK967010.1 | Pseudomonadota | Gammaproteobacteria | Pseudomonadales | Pseudomonadaceae | *A. aurita* medusa Baltic Sea | Marine Bouillon | 30°C | enrichment/ first screening/ host range |
| 62 | *Sulfitobacter pontiacus* | OQ398158 | Pseudomonadota | Alphaproteobacteria | Rhodobacterales | Rhodobacteraceae | *M. leidy* Baltic Sea | Marine Bouillon | 30°C | host range |
| 97 | *Shewanella sp.* | OQ398161 | Pseudomonadota | Gammaproteobacteria | Alteromonadales | Shewanellaceae | *A. aurita* polyp North Sea husbandry | Marine Bouillon | 30°C | host range |
| 199 | *Staphylococcus aureus* | OQ398168 | Bacillota | Bacilli | Bacillales | Staphylococcaceae | Artificial Seawater 30 PSU | Marine Bouillon | 30°C | host range |
| DSMZ 11823 | *Staphylococcus aureus* | DSMZ 11823 | Bacillota | Bacilli | Bacillales | Staphylococcaceae | clinical material | Trypticase Soy Yeast Broth | 37°C | host range |
| 67 | *Staphylococcus aureus* | OQ398159 | Bacillota | Bacilli | Bacillales | Staphylococcaceae | *M. leidy* Baltic Sea husbandry | Marine Bouillon | 30°C | host range |
| 102 | *Staphylococcus aureus* | OQ398162 | Bacillota | Bacilli | Bacillales | Staphylococcaceae | *A. aurita* polyp North Sea husbandry | Marine Bouillon | 30°C | host range |
| 158 | *Staphylococcus aureus* | OQ398165 | Bacillota | Bacilli | Bacillales | Staphylococcaceae | Artificial Seawater 18 PSU | Marine Bouillon | 30°C | host range |
| 161 | *Staphylococcus aureus* | OQ398166 | Bacillota | Bacilli | Bacillales | Staphylococcaceae | Artificial Seawater 18 PSU | Marine Bouillon | 30°C | host range |
| 127 | *Staphylococcus aureus* | OQ398163 | Bacillota | Bacilli | Bacillales | Staphylococcaceae | *A. aurita* polyp North Atlantic husbandry | Marine Bouillon | 30°C | host range |
| DSMZ 28319 | *Staphylococcus epidermidis* | DSMZ 28319 | Bacillota | Bacilli | Bacillales | Staphylococcaceae | catheter sepsis | Trypticase Soy Yeast Broth | 37°C | host range |
| DSMZ 20328 | *Staphylococcus hominis* | DSMZ 20328 | Bacillota | Bacilli | Bacillales | Staphylococcaceae | human skin | Trypticase Soy Yeast Broth | 37°C | host range |
| DSMZ 100616 | *Staphylococcus warneri* | DSMZ 100616 | Bacillota | Bacilli | Bacillales | Staphylococcaceae | cleanroom facility, TAS | Trypticase Soy Yeast Broth | 30°C | host range |
| DSMZ 12643 | *Streptococcus mitis* | DSMZ 12643 | Bacillota | Bacilli | Lactobacillales | Streptococcaceae | oral cavity, human | Trypticase Soy Yeast Broth | 37°C | host range |
| DSMZ 20523 | *Streptococcus mutans* | DSMZ 20523 | Bacillota | Bacilli | Lactobacillales | Streptococcaceae | carious dentine | Trypticase Soy Yeast Broth | 37°C | host range |
| 296 | *Citrobacter braakii* | OQ398170 | Pseudomonadota | Gammaproteobacteria | Enterobacterales | Enterobacteriaceae | *A. aurita* polyp North Atlantic husbandry | Marine Bouillon | 30°C | host range |
| 283 | *Citrobacter freundii* | OQ398169 | Pseudomonadota | Gammaproteobacteria | Enterobacterales | Enterobacteriaceae | *A. aurita* polyp Baltic Sea husbandry | Marine Bouillon | 30°C | host range |
| 321 | *Citrobacter freundii* | OQ398172 | Pseudomonadota | Gammaproteobacteria | Enterobacterales | Enterobacteriaceae | Artificial Seawater 30 PSU | Marine Bouillon | 30°C | host range |
| 313 | *Citrobacter* sp. | OQ398171 | Pseudomonadota | Gammaproteobacteria | Enterobacterales | Enterobacteriaceae | Artificial Seawater 18 PSU | Marine Bouillon | 30°C | host range |
| DSMZ 18039 | *Escherichia coli* | DSMZ 18039 | Pseudomonadota | Gammaproteobacteria | Enterobacterales | Enterobacteriaceae | unknown source | Luria-Bertani Bouillon | 37°C | host range |
| strain 8 | *Escherichia coli* | (Hanahan, 1983) | Pseudomonadota | Gammaproteobacteria | Enterobacterales | Enterobacteriaceae | unknown source | Luria-Bertani Bouillon | 37°C | host range |
| **DSMZ 30083** | *Escherichia coli* | DSMZ 30083 | Pseudomonadota | Gammaproteobacteria | Enterobacterales | Enterobacteriaceae | urine | Luria-Bertani Bouillon | 37°C | host range |
| **DSMZ 13698** | *Escherichia fergusonii* | DSMZ 13698 | Pseudomonadota | Gammaproteobacteria | Enterobacterales | Enterobacteriaceae | faeces of 1-year-old boy | Luria-Bertani Bouillon | 37°C | host range |
| **strain 27** | *Klebsiella oxytoca* | Prof. Dr. Podschun, (National Reference Laboratory for Klebsiella species, Kiel University) | Pseudomonadota | Gammaproteobacteria | Enterobacterales | Enterobacteriaceae | unknown source | Nutrient Broth | 30°C | host range |
| **DSMZ 30104** | *Klebsiella pneumoniae* | DSMZ 30104 | Pseudomonadota | Gammaproteobacteria | Enterobacterales | Enterobacteriaceae | unknown source | Nutrient Broth | 30°C | host range |
| **DSMZ 4782** | *Shigella flexeneri* | DSMZ 4782 | Pseudomonadota | Gammaproteobacteria | Enterobacterales | Enterobacteriaceae | unknown source | Caso Bouillon | 37°C | host range |
| **DSMZ 1707** | *Pseudomonas aeruginosa* | DSMZ 1707 | Pseudomonadota | Gammaproteobacteria | Pseudomonadales | Pseudomonadaceae | unknown source | Caso Bouillon | 30°C | host range |
| **9** | *Pseudomonas sp.* | OQ398155 | Pseudomonadota | Gammaproteobacteria | Pseudomonadales | Pseudomonadaceae | *A. aurita* medusa Baltic Sea | Marine Bouillon | 30°C | host range |
| **170** | *Pseudomonas sp.* | OQ398167 | Pseudomonadota | Gammaproteobacteria | Pseudomonadales | Pseudomonadaceae | Artificial Seawater 18 PSU | Marine Bouillon | 30°C | host range |
| **DSMZ 50256** | *Pseudomonas syringae* | DSMZ 50256 | Pseudomonadota | Gammaproteobacteria | Pseudomonadales | Pseudomonadaceae | Triticum aestivum, glume rot of wheat | Caso Bouillon | 30°C | host range |
| **24** | *Maribacter sp.* | MK967024.1 | Bacteroidota | Flavobacteriia | Flavobacteriales | Flavobacteriaceae | *A. aurita* medusa Baltic Sea husbandry | Marine Bouillon | 30°C | host range |
| **79** | *Olleya sp.* | MK967053.1 | Bacteroidota | Flavobacteriia | Flavobacteriales | Flavobacteriaceae | *A. aurita* polyp Baltic Sea husbandry | Marine Bouillon | 30°C | host range |
| **98** | *Olleya marilimosa* | MK967072.1 | Bacteroidota | Flavobacteriia | Flavobacteriales | Flavobacteriaceae | *A. aurita* polyp North Sea husbandry | Marine Bouillon | 30°C | host range |
| **108** | *Chryseobacterium hominis* | MK967082.1 | Bacteroidota | Flavobacteriia | Flavobacteriales | Flavobacteriaceae | *A. aurita* polyp North Sea husbandry | Marine Bouillon | 30°C | host range |
| 57 | *Olleya marilimosa* | MK967032.1 | Bacteroidota | Flavobacteriia | Flavobacteriales | Flavobacteriaceae | *M. leidy*<br>Baltic Sea | Marine Bouillon | 30°C | host range |
| 199 | *Maribacter sp.* | MK967170.1 | Bacteroidota | Flavobacteriia | Flavobacteriales | Flavobacteriaceae | Artificial<br>Seawater 30<br>PSU | Marine Bouillon | 30°C | host range |

### Taxonomic classification of bacterial isolates

Bacteria were enriched and isolated from *A. aurita* medusae collected in the Baltic Sea, as described in the previous study by Weiland-Bräuer *et al*., 2020 (Weiland-Bräuer et al., 2020). Additional isolates previously not published were taxonomically classified in the present study. The bacterial isolates were grown, and genomic DNA was isolated from overnight cultures (5 mL) using the Wizard Genomic DNA Purification Kit (Promega GmbH, Walldorf) according to the manufacturer’s instructions. 16S rRNA genes were PCR-amplified from 50 ng isolated genomic DNA using the bacterium-specific 16S rRNA gene primer 27F (5′-AGAGTTTGATCCTGGCTCAG-3′) and the universal primer 1492R (5′- GGTTACCTTGTTACGACTT-3′) (Zavaleta et al., 1996) resulting in a 1.5 kb PCR fragment. The fragments were Sanger sequenced at the sequencing facility at the Institute of Clinical Molecular Biology, University of Kiel (IKMB). Sequence analysis was conducted using CodonCode Aligner v. 9.0. Sequence data of full-length 16S rRNA genes are deposited under GenBank accession numbers OQ397638- OQ397653 and OQ398153-OQ398172.

### Phage enrichment, isolation, and purification

Water columns were sampled in the Baltic Sea at Kiel fjord (54.329649, 10.149129) in March 2020 and June 2021. Samples were taken from the surface (< 50 cm depth) using a sterile 20 L canister. 50 mL of the seawater samples and 50 mL of MB medium were mixed with 1 mL of an overnight culture of a mixture of potential host bacterial strains (**Table 1**, column [use in the study], category [enrichment/first screening], a total of 55 strains) and placed on a shaking incubator (120 rpm) at 30°C for 24 h. The mixture was centrifuged at 4,000 × *g* for 30 min, and the supernatant passed through a 0.22 μm pore-size polycarbonate syringe filter (Sartorius, Goettingen) to remove the residual bacterial cells.

The spot test assay, a procedure based on the double-layer plaque technique (Cormier & Janes, 2014), was used as an initial test to detect lytic phages by plaques on all 55 bacterial strains, separately grown in top-agar on agar plates. Briefly, 10 µL of each filtered mixture was spotted on an MB 1.5 % agar plate containing a second solidified layer of 3 mL 0.6 % MB top agar mixed with 100 µL of a single bacterial host strain. The plates were incubated overnight at 30 °C. Plaques generated by bacteriophage-induced bacterial lysis were detected the following day. Plaques were exclusively detected for bacterial isolates No. 8 (*Pseudomonas sp*. MK967010.1), No. 7 (*Citrobacter sp*., OQ398154), and No. 6 (*C. freundii*, OQ398153). Those three bacterial isolates were used as bacterial host strains for the following assays to study phage characteristics.

### Phage purification, titration, and propagation

Original phage plaques were used for further purification of phages. Morphologically distinct plaques were picked from the agar plate using a sterile toothpick and streaked on a freshly prepared double-agar plate with the respective host strain in the top agar. The procedure was repeated three times to ensure pure and single phages. Phage lysate preparations were conducted from top agar plates with approx. 10^5^ plaque-forming units per mL (pfu/mL). Phage-containing top agar was collected with a sterile loop and transferred into a 15 mL Falcon tube (Sarstedt, Germany). 3 mL of liquid MB was added. The mixture was vortexed and centrifuged at 4,000 × *g* for 10 min. The supernatant was filtered through a 0.22 μm polycarbonate syringe filter (Sartorius, Goettingen) to remove bacterial cells and agar debris. The respective pfu/mL of resulting phage lysate was determined, and the lysate was stored at 4 °C. Phage stability at 4 °C was analyzed every 2 days for 2 weeks using the double-layer agar method with no significant variance in pfu/mL.

Phage propagation was performed in liquid culture. 1 mL of the respective bacterial host (overnight culture) was inoculated into 48 mL of MB in a 100 mL Erlenmeyer flask and incubated at 30 °C and 120 rpm until OD_600nm_ = 0.2 - 0.3 was reached. 1 mL of the phage lysate (10^6^ pfu/mL) was added to the cultures, which were further incubated for 3 h at 30 °C with shaking. The culture was transferred in a 50 mL Falcon tube (Sarstedt, Germany) and centrifuged at 4,000 × *g* for 10 min. The supernatant was sterile-filtered using a 0.22 μm syringe filter (Sartorius, Goettingen). Phage lysates were stored at 4 °C.

### Transmission electron microscopy

50 mL of freshly prepared phage lysate (> 10^8^ pfu/mL) was ultracentrifuged (Optima XE-100 ultracentrifuge, Beckman Coulter, Brea, CA, USA) at 109, 800 × *g* for 30 min. Phage pellets were resuspended in 1 mL of Ultra-pure water (Carl Roth, Karlsruhe, Germany) overnight at 4 °C on a 3D shaker. Subsequently, TEM grids (copper, 400 square mesh, formvar-coated) were glow-discharged for 60 s at 0.6 mbar air pressure, and 10 mA glow current using a Safematic CCU-010 unit and then incubated with 8 μL of the phage lysate (> 10^8^ pfu/mL) for 5 min. Grids were washed briefly on six drops of water, stained with 1 % uranyl acetate for 10 s, blotted to remove the excess stain, and air dried. Samples were imaged with a Tecnai G2 Spirit BioTwin transmission electron microscope operated with a LaB6 filament at 80 kV, and equipped with an Eagle 4k HS CCD camera, TEM User interface (v. 4.2), and TIA software (v. 2.5) (all FEI / Thermo Fisher Scientific). Open-source Fiji software was used to measure the head width (perpendicular to the vertical axis) and the tail length of phages.

### One-step growth curve analysis

Bacterial host strains were grown overnight in 5 mL MB. 500 µL of the overnight culture were incubated in 50 mL MB at 30 °C until turbidity at 600 nm of T_600_ = 0.1 - 0.2 (10^7^ cells/mL) were reached. 10 mL of the bacterial culture were centrifuged at 4,000 x *g* for 10 min at 4 °C. The cell pellets were resuspended in 5 mL MB medium, and 5 mL of phage lysate (10^7^ pfu/mL) was added with a multiplicity of infection (MOI) of 1.0. Phages were allowed to adsorb to the bacterial host cells during 5 min incubation at 30 °C. Afterwards, the mixture was centrifuged at 4,000 × *g* for 10 min to remove the free phage particles before resuspending the samples in 10 mL MB. The phage titer was immediately determined by double-agar layer plaque assay (t_0_). The mixture was incubated at 30 °C, aliquots were collected in 10 min intervals over a 120 min period, and phage titers were determined. Three independent experiments were performed for each phage.

### Efficiency of plating and host range

The host range of isolated bacteriophages was initially determined by the spot assay and verified by the double-layer agar method. A selection of 43 strains belonging to different genera (**Table 1**, column [use in the study], category [host range]) was tested. The bacterial strains were individually grown overnight in 5 mL cultures. An aliquot of 100 μL of each culture and 3 mL of the respective culture medium containing 0.6 % agar was mixed and poured onto an agar plate. After 15 min at room temperature, to allow the top agar to solidify, 10 μL of the 10-fold serially diluted phage lysate (original 10^9^ PFU/mL, diluted in MB) were spotted onto the soft agar. The plates were then incubated at the respective incubation temperature of the host strain (**Table 1**). Plaques were examined after 16 h of incubation. The Efficiency of Plating (EOP) was calculated by the ratio of the average PFU on a tested host to the average PFU on a corresponding reference (original) host. The variation of EOP values is represented as a heat map using Excel.

### Viral DNA isolation

Genomic DNA was isolated from the phage lysates using a modified phenol-chloroform-isoamyl alcohol method (Nale et al., 2016). Briefly, 500 µL of phage lysates (10^12^ PFU/mL) were treated with 1 U/ml of DNase I and 1 U/ml of RNase A (Thermo Fisher, Germany) in a Reaction Buffer (100 mM Tris-HCl, 25 mM MgCl_2_, 1 mM CaCl_2_) (Thermo Fisher, Germany), and incubated 1 h at 37 °C to remove external nucleic acids. Afterwards, 0.5 M EDTA, 40 µL Proteinase K (20 mg/mL, Thermo Fisher, Germany), 1 M CaCl_2,_ and 20 % sodium dodecyl sulfate (Roth, Germany) were added before incubating at 37 °C for 2 h. Following, samples were incubated for a further 2 h at 65 °C. Viral DNA was extracted with an equal volume (vol) of phenol-chloroform-isoamyl alcohol (25:24:1, Roth, Germany) and centrifuged at 3,000 × *g* for 15 min. The step was repeated, and the supernatant was transferred to phase lock tubes (Quantabio, Germany). The aqueous phase was mixed with an equal volume of chloroform and centrifuged at 1,600 x *g* for 15 min. The aqueous layer was mixed with 3 M sodium acetate and 1 vol of isopropanol to precipitate the DNA. After incubation overnight at -20 °C, the DNA was pelleted by centrifugation at 12,600 × *g* for 30 min at 4 °C. The pellet was washed with 70 % ethanol, air-dried, and dissolved in Ultra-pure water (Roth, Germany). DNA was stored at -20 °C before sequencing. DNA quality and quantity were analyzed using a NanoDrop1000 spectrophotometer and a Qubit double-stranded BR assay kit on a Qubit fluorometer (Thermo Fisher, Germany).

### Sequencing, bioinformatic analysis, and annotation of viral genomes

Long sequencing reads were obtained using the Oxford Nanopore Technologies MinION platform (R9.4.1 flow cell). The MinION sequencing library was prepared according to the manufacturer’s guidelines using the SQK-RBK004 Rapid Barcoding Kit. MinION sequencing was performed with MinKNOW v. 22.08.4. The raw sequencing data (fast5 format) were base-called using Guppy v. 6.2.7, and finally, demultiplexing was performed using qcat v.1.1.0. Quality assessment and adapter trimming of the MinION long-read sequences of the four viral genomes was performed using LongQC v1.2.0 (Fukasawa et al., 2020) and Filtlong v.0.2.1 (https://github.com/rrwick/Filtlong). Filtered sequence reads with an average length > 1,000 bps were selected, omitting the worst 5 % of reads. Reads per genome were assembled using the assembler Flye v2.9 (Kolmogorov et al., 2019), resulting in complete, single-contig genomes. The number of sequenced reads before and after filtration, the GC content, reads coverage, and N50 values are provided in **Table 2**. The completeness and contamination of the assembled viral genomes were assessed using CheckV (Nayfach et al., 2021).

**Table 2.** Viral genome characteristics and overview of assembly-related metrics.

| Phage | NCBI<br>Accession<br>No. | No.<br>of<br>reads | No. of<br>filtered<br>reads | Sequence<br>coverage | N50 | Genome<br>length<br>(bps) | GC<br>content<br>(%) | predicted<br>ORFs | unknown<br>proteins |
| --- | --- | --- | --- | --- | --- | --- | --- | --- | --- |
| *Pseudomonas*<br>phage BSwM<br>KMM1 | OP902294 | 4.085 | 3.214 | 247.488 | 17.553 | 137.386 | 31.77 | 259 | 200 |
| *Citrobacter* phage<br>BSwM KMM2 | OP902295 | 810 | 595 | 74.163 | 22.118 | 88.537 | 39.55 | 137 | 94 |
| *Citrobacter* phage<br>BSwS KMM3 | OP902292 | 837 | 598 | 130.676 | 20.517 | 49.164 | 43.17 | 92 | 58 |
| *Citrobacter* phage<br>BSwM KMM4 | OP902293 | 6.433 | 5.371 | 544.327 | 23.894 | 86.911 | 39.02 | 138 | 100 |

Viral genomes were annotated using Prokka v.1.14.6 (Seemann, 2014) and the “--kindgom Viruses” option. Functional annotation of the four isolated phages was performed using EggNOG mapper v2.1.9 (Cantalapiedra et al., 2021) and the eggNOG database v5.0 (Huerta-Cepas et al., 2019), and the sequence searches were performed using DIAMOND (Buchfink et al., 2021). Putative Auxiliary metabolic genes were additionally annotated (AMG) using DRAM-v with default databases (Shaffer et al., 2020), the viral mode of DRAM (Distilled and Refined Annotation of Metabolism). For this purpose, the four isolated genomes were processed through Virsorter2 v2.2.4 (Guo et al., 2021) with the “-- prep-for-dramv” parameter to generate an “affi-contigs.tab” needed by DRAM-v to identify AMGs. Putative AMGs were identified based on the resulting assigned auxiliary score (AMG score <=3) and metabolic flag (M flag, no V flag, no A flag). COG (Galperin et al., 2015) and KEGG (Kanehisa et al., 2007) annotations were derived from the EggNOG mapper and DRAM-V results. vConTACT v2.0 (Bin Jang et al., 2019) with default settings were used to cluster the four viral genomes together with the sequences from the “ProkaryoticViralRefSeq207-Merged” to generate Viral Clusters (VCs) and determine the genus-level taxonomy of the viral genomes.

Relationships between the four isolated phages and reference genomes were analyzed using nucleic acid-based intergenomic similarities calculated with VIRIDIC (Virus Intergenomic Distance Calculator) v1.0r3.6 using default settings (Moraru et al., 2020). VIRIDIC identifies intergenomic nucleotide similarities between viruses using BLASTN pairwise comparisons and organizes viruses into clusters (genera ≥ 70 % similarities and species ≥ 95 % similarities). These cut-offs assign viruses into ranks following the International Committee on Taxonomy of Viruses (ICTV) genome identity thresholds. The reference genomes were selected based on the results of the gene-sharing network analysis where the four isolated phages clustered with viruses from the Prokaryotic Viral RefSeq Database. Genome-based phylogeny and classification of the four isolated viruses together with the same reference prokaryotic viruses were performed using the VICTOR web service (Virus Classification and Tree Building Online Resource). VICTOR is a Genome-BLAST Distance Phylogeny (GBDP) method that computes pairwise comparisons of the amino acid sequences (including 100 pseudo-bootstrap replicates) and uses them to infer a balanced minimum evolution tree with branch support via FASTME, including subtree pruning and regrafting postprocessing (Lefort et al., 2015) for each of the formulas D0, D4, and D6, respectively. Trees were rooted at the midpoint (Farris, 1972) and visualized with ggtree (Yu, 2020). The OPTSIL algorithm (Göker et al., 2009), the suggested clustering thresholds (Meier-Kolthoff & Göker, 2017), and an F-value (fraction of linkages necessary for cluster fusion) of 0.5 were used to estimate taxon boundaries at the species, genus, and family levels for prokaryotic viruses (Meier-Kolthoff et al., 2014). The position and annotation of predicted viral genes in the phage genomes were visualized using Clinker v0.0.27 (Gilchrist & Chooi, 2021). Isolated viruses were compared and visualized to the closest related species determined based on intergenomic similarity analysis. Clinker generates global alignments of amino acid sequences based on the BLOSUM62 substitution matrix. A 0.5 identity threshold was used to display the alignments. The complete phage genome sequences (assemblies) are available at NCBI under the accession numbers OP902292- OP902295. Raw sequence reads were deposited on the Sequence Read Archive (SRA) under BioProject PRJNA908753 and accession numbers SRR22580853, SRR22580850, SRR22580849, and SRR22580845.

## Results

Bacteriophages were isolated from the Baltic Sea water. Four novel phages infecting bacterial colonizers of the Cnidarian moon jellyfish *A. aurita* were identified and characterized.

### Isolation of bacteriophages from Baltic seawater targeting marine bacteria associated with A. aurita

Seawater samples from the Kiel fjord (Baltic Sea) were used for phage enrichments with 55 bacteria associated with *A. aurita*. The bacteria chosen were previously described (Weiland-Bräuer et al., 2020) and represent a diverse set of abundant species associated with *A. aurita,* possessing varying forms, colors, and colony morphologies (**Table 1**, (Weiland-Bräuer et al., 2015)). One lytic phage targeting *Pseudomonas sp*. (isolate No. 8, MK967010.1), one targeting *Citrobacter freundii* (isolate No. 6, OQ398153), and two phages targeting *Citrobacter sp*. (isolate No. 7, OQ398154) were isolated. Those bacteriophages were designated as *Pseudomonas* phage BSwM KMM1, *Citrobacter* phage BSwM KMM2, *Citrobacter* phages BSwS KMM3, and *Citrobacter* phages BSwM KMM4. In the following, phage designations are abbreviated to KMM1 – KMM4.

### Plaque and virion morphology assign the phages to the class of Caudoviricetes

The identified lytic bacteriophages KMM1 (*Pseudomonas* phage), KMM2 (*Citrobacter freundii* phage), KMM3 and KMM4 (*Citrobacter sp.* phages) formed clear plaques with well-defined boundaries when infecting the respective host bacterial strain after 16 h of incubation at 30 °C. Notably, infection with KMM3 resulted in a clear center and a turbid surrounding halo (**Fig. 1A** and **Table 3**). Lysis plaques were further differentiated by the halo size. Phages KMM1, KMM2, and KMM4 generated plaques with a diameter varying between 0.8 mm and 1.2 mm, while KMM3 showed plaques with a diameter of 3 mm (**Fig. 1A** and **Table 3**).

**Figure 1.**
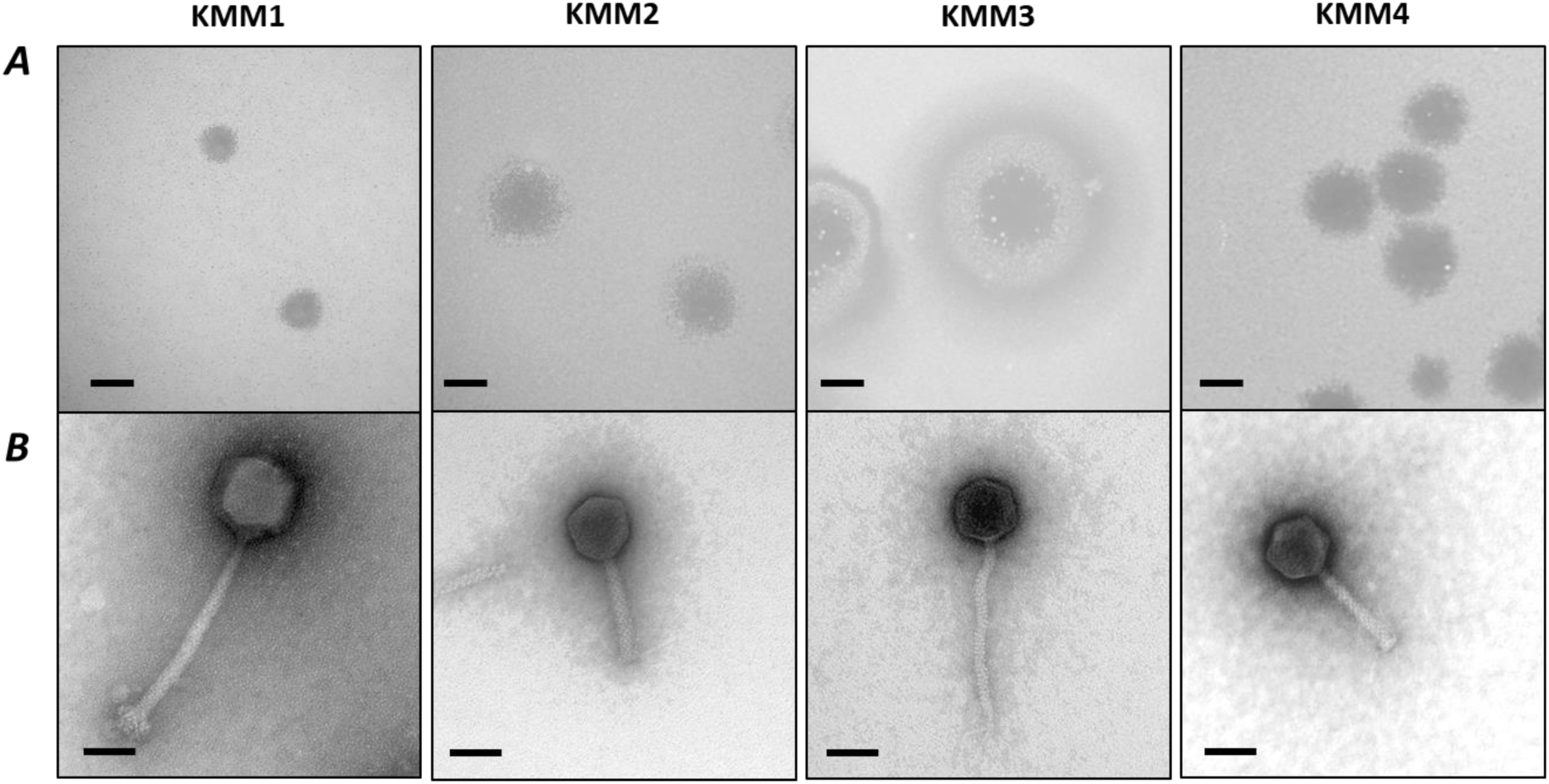
Plaque and virion morphology of isolated bacteriophages KMM1-KMM4. (***A***) Plaque morphologies were detected on MB double-agar layer plates after 16 h of incubation at 30 °C. Plaques formed on a lawn of: *Pseudomonas sp.,* MK967010.1 (**KMM1**), *Citrobacter freundii,* OQ398153 (**KMM2**), and *Citrobacter sp.,* OQ398154 (**KMM3**, **KMM4**). Scale bars represent 1 mm. (***B***) Transmission electron micrographs of phage lysates KMM1 – KMM4, scale bars represent 50 nm.

**Table 3.** Characteristics of isolated bacteriophages KMM1 – KMM4. Plaque properties (N = 10), phage morphology (tail, N= 20; head, N = 10), latent period, and burst size (N = 3) are listed.

|  | KMM1 | KMM2 | KMM3 | KMM4 |
| --- | --- | --- | --- | --- |
| **Plaques properties** | Small, round, and clear<br>Size: 0.8-1 mm | Small, round, and clear<br>Size: 1-1.5 mm | Big, round with a clear center and surrounding turbid halo<br>Size: 2.5-3.5 mm | Small, round, and clear<br>Size: 1.2-1.5 mm |
| **Phage morphology** |  |  |  |  |
| Tail length (nm) | 176.4 ± 8.4 | 90.9 ± 3.9 | 131.7 ± 9.3 | 90.5 ± 4.5 |
| Head width (nm) | 75.8 ± 3 | 58.2 ± 2.7 | 50 ± 1.9 | 57.5 ± 2.8 |
| Head shape | icosahedral | icosahedral | icosahedral | icosahedral |
| **Latent period (min)** | 27 | 20 | 45 | 30 |
| **Burst size (phages/infected bacterial cell)** | 55 | 280 | 60 | 120 |
| **Taxonomy prediction** | *Myovirus* | *Myovirus* | *Siphovirus* | *Myovirus* |

Transmission Electron Microscopy (TEM) images revealed a pre-classification of all phages to the class of *Caudoviricetes*, characterized by long tails with a collar, base plates with short spikes, six long kinked tail fibers, and isometric heads (**Fig. 1**). Imaging further indicated that KMM1, KMM2, and KMM4 could be assigned to the *Myovirus* family by possessing contractile tails, while KMM3 can be classified to the *Siphovirus* family due to a non-contractile tail (**Fig. 1** and **Table 3**). The width of the phage heads of KMM2, KMM3, and KMM4 ranged from 50 ± 1.9 nm to 58.2 ± 2.7 nm. The tail length was similar for KMM2 and KMM4, with an average length of 90.7 ± 4.2 nm, while the tail length of KMM3 was 131.7 ± 9.3 nm. Phage KMM1 was the largest of the isolated phages, with a head width of 75.8 ± 3 nm and a tail length of 176.4 ± 8.4 nm (**Table 3**).

### Phages KMM1, KMM2, and KMM4 have a shorter lytic cycle than phage KMM3

To assess each phage’s capacity for infection, one-step growth curves were conducted with the respective host strains in MB medium at 30 °C for 120 min in three independent biological replicates (**Fig. 2**). The calculated values for latent time and burst size are displayed in **Table 3**. KMM1 and KMM2 each showed an approximately 20 min latent period resulting in a burst size of 55 pfu/cell after 100 min (KMM1), while KMM2 released an average yield of 280 pfu/cell after 110 min. KMM4 infection resulted in a prolonged latent period of 30 min leading to 120 released phages (pfu/cell) after 100 min. KMM3 showed the most extended latent period with 45 min and was characterized by a burst size of 60 pfu/cell, reached after 90 min.

**Figure 2.**
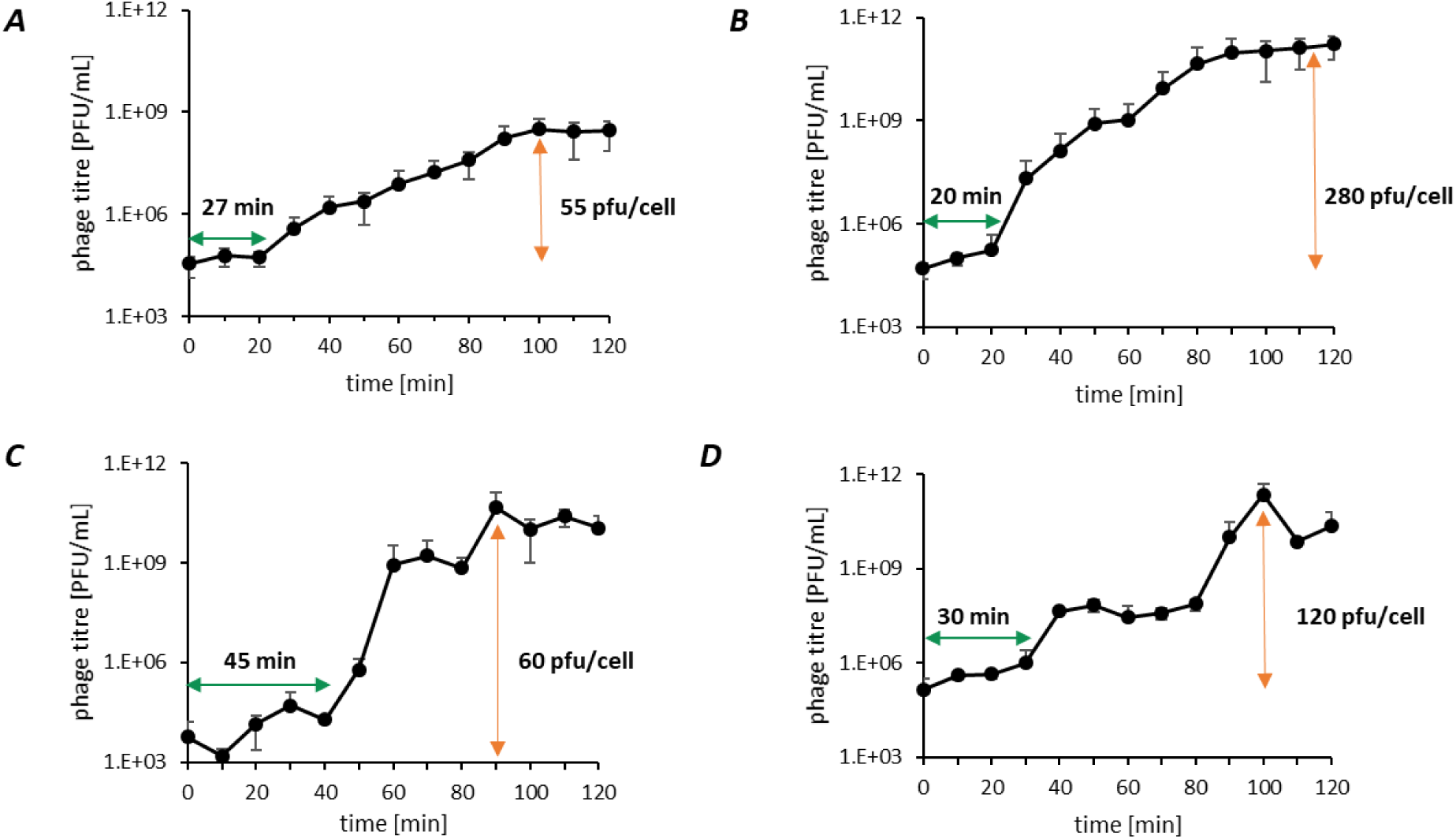
Infection cycles of isolated phages. One-step growth curves over 120 min were performed to calculate the latent period (green arrow) and burst size (orange arrow). (***A***) KMM1 infected *Pseudomonas sp.* (MK967010.1) after 27 min with the release of 55 pfu/cell, (**B**) KMM2 infected *Citrobacter freundii* (OQ398153) after 20 min with the release of 280 pfu/cell, (**C**) KMM3 and (**D**) KMM4 infected *Citrobacter sp.* (OQ398154) after 45 and 30 min, respectively, with the release of 60 and 120 pfu/cell, respectively. Values represent the mean of three biological replicates.

### KMM1 is a powerful broad-host-range phage, whereas KMM2, KMM3, and KMM4 are narrow-host-range phages

The KMM1 phage was initially found to infect *Pseudomonas sp.*, while KMM2, KMM3, and KMM4 were shown to infect *Citrobacter spp.*, which are phylogenetically classified in the *Pseudomonadaceae* and *Enterobacteriaceae*, respectively. The host range of the phages was determined by spot assays on 43 strains (**Table 1**, column [use in the study], category [host range]) belonging to the same genera, *Pseudomonas* and *Citrobacter*.

Furthermore, phages were tested against representatives of *Shewanellaceae* and *Rhodobacteraceae* of phylum Proteobacteria, *Staphylococcaceae* and *Streptococcaceae* of Bacilli, and *Chryseobacterium, Olleya,* and *Maribacter* of the abundant class of Flavobacteriia present in the *A. aurita*-associated microbiota. Bacterial sensitivity to a given bacteriophage was evaluated based on the occurrence of a lysis halo. Additionally, the respective phage efficiency of plating (EOP) was determined with those bacteria showing lysis in the spot tests. EOP for each host bacterium was calculated by comparing it with a score of 10^9^ pfu/mL obtained for the original host infection. As shown by the heatmap in **Fig. 3**, KMM1 infects, in addition to the primary host, two strains of *Pseudomonas*, one *Shewanella* strain also belonging to Gamma-Proteobacteria, one *Sulfitobacter* strain belonging to Alpha-Proteobacteria, and 14 strains of the Gram-positive family *Staphylococcaceae*, two of which with a slightly higher EOP. The phages KMM2, KMM3, and KMM4 showed comparable and narrow host ranges within the genus *Citrobacter*. However, the titers observed for the phages were different and thus the EOP, as shown by the value-based color code (**Fig. 3**). The phages KMM2 and KMM4 were further able to infect the *Enterobacteriaceae* bacterium *Shigella flexneri*. In contrast, the phage KMM3 infected two *Escherichia coli* strains of *Enterobacteriaceae*. The phages infected none of the Flavobacteriia representatives.

**Figure 3.**
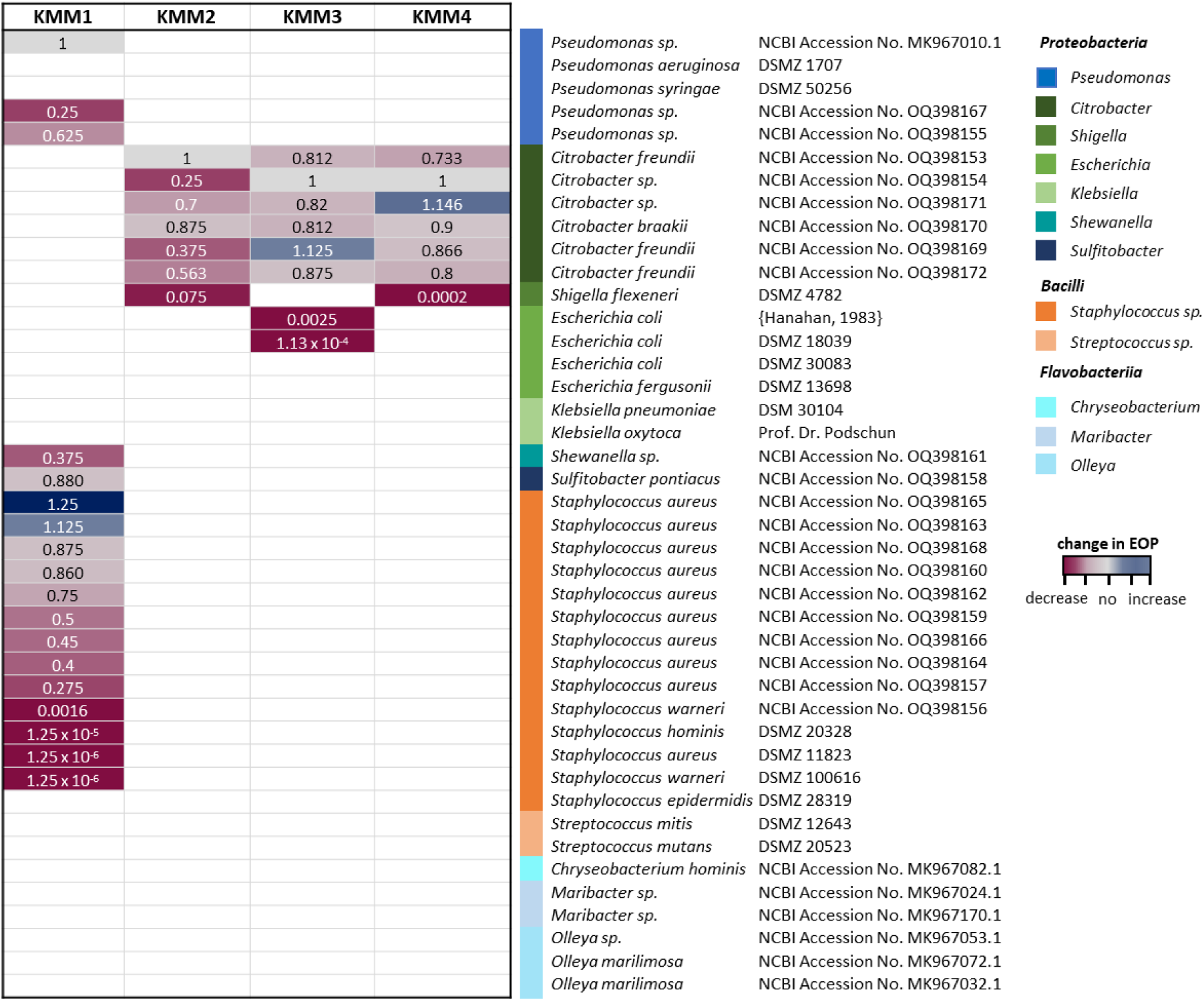
Host range of isolated phages. Phages KMM1 – KMM4 were used for infection assays with selected taxons (color-code on the right categorizes taxons into classes). The efficiency of plating (EOP) for each host bacterium was calculated by comparing it with a score of 109 pfu/mL for the original host infection (value = 1). Missing coloring indicates no infection.

### Novel phage species confirmed by genome sequencing analysis

The viral genomes of the highly effective lytic phages were sequenced using Nanopore technology. Complete phage genomes were assembled (NCBI Accession Nos. OP902292-OP902295) from Nanopore long reads of the double-stranded DNA. **Table 2** summarizes the key information regarding sequencing, assembly, and annotation. Three of the four viral genomes were assigned “high-quality” (> 90 % completeness), while phage KMM2 was assigned as “complete” due to the presence of direct terminal repeats (DTR), which may indicate a circular genome. KMM1 has a 137 kbp genome with a GC content of 31%, while KMM2 and KMM4 showed genome sizes of approx. 87 kbp bp with an average GC content of 39 %. The KMM3 genome was found to be the smallest, with 49 kbp but with the highest GC content of 43 %. In total, 259 putative ORFs were predicted in the genome of phage KMM1, 137 and 138 ORFs were predicted in KMM2 and KMM4 genomes, respectively, and 92 ORFs in the KMM3 genome (**Table 2**). Phages were clustered into species and higher-order groups to investigate phage genomic diversity and identify closely related groups of phages. Viral Clusters (VCs) and genus-level taxonomy of the four isolated and sequenced viral genomes were generated. VICTOR (based on pairwise whole genome distance comparisons) was used to compare 96 previously described viral taxa with the phage genomes of KMM1 – KMM4. The results indicated that the four identified phages belong to three different genera within the *Caudovirales* class, as they are grouped into three different clades in the phylogenetic tree (**Fig. 4**). The four phages are members of “*Heunggongvirae*; *Uroviricota*; *Caudoviricetes*; *Caudovirales*” and more specifically KMM1 is a member of the *Herelleviridae* family, KMM2 and KMM4 are members of “*Caudovirales*; *Myoviridae*; *Mooglevirus*; *Felixounavirus*” and *Suspvirus* genus, and KMM3 is classified as a member of “*Caudovirales*; *Drexlerviridae*; *Tlsvirus*” and *Hicfunavirus* genus. Based on VICTOR and vConTACT2 analysis, the four isolated phages belong to four predicted genera and three species (**Fig 4**). VIRIDIC (Virus Intergenomic Distance Calculator) was used to determine the pairwise intergenomic similarities between the phage genomes characterized in this work compared to reference genomes. The intergenomic analysis gave evidence that the isolated viral genomes are four novel species.

**Figure 4.**
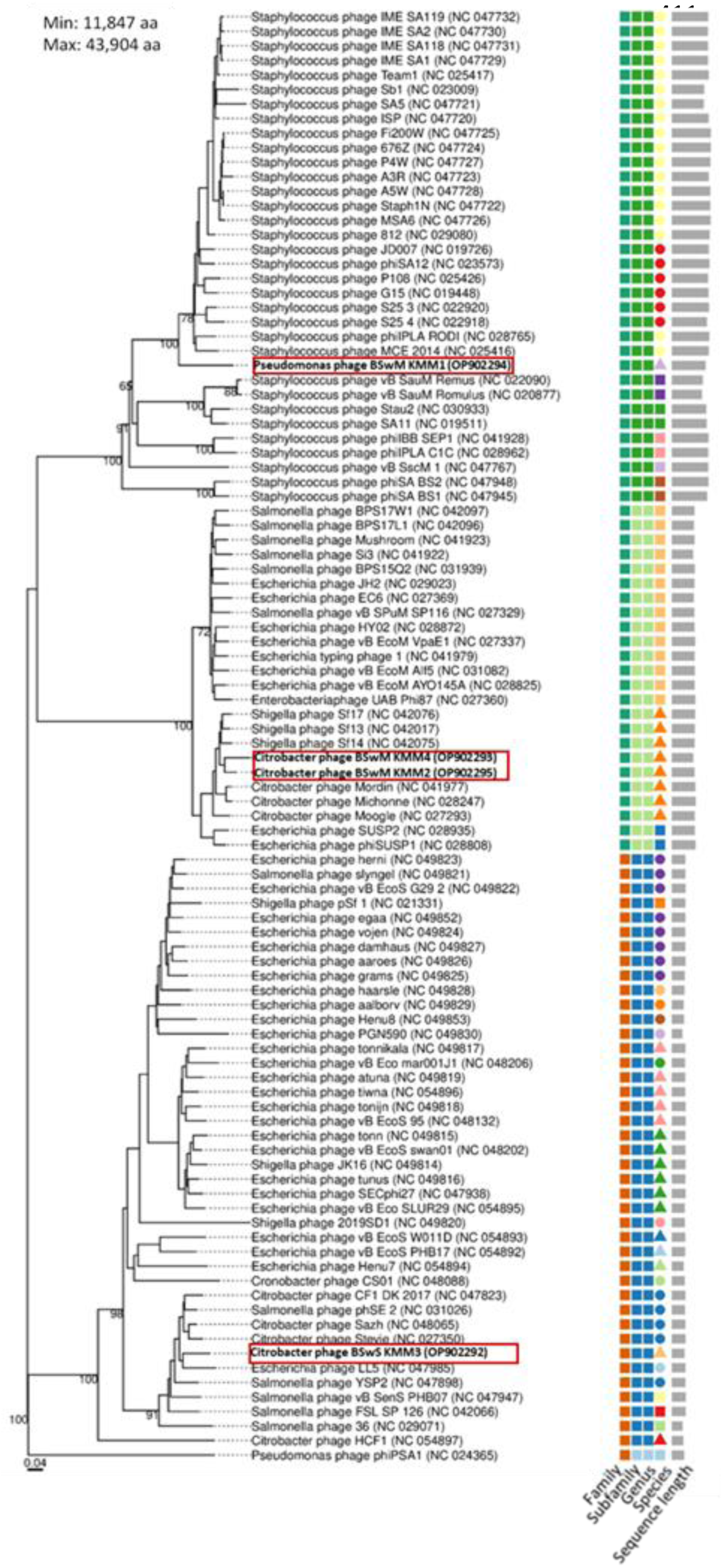
Taxonomic classification of phages KMM1 – KMM4. Phylogenetic tree of isolated phages KMM1 – KMM4 (red rectangles) was generated with the whole genome-based VICTOR analysis. Phages belonging to different families, subfamilies, genera, and species were color coded. The scale represents homology in %.

Although experimental results revealed four different phages, genome analyses using VICTOR and VIRIDIC resulted in contrasting statements. Both programs confirmed that KMM1 and KMM3 are novel species. However, those analyses did not resolve whether KMM2 and KMM4 were one or two species or whether they were novel. In the next step, genomes were annotated and compared to their best homologs (**Fig. 5**). Open reading frames (ORFs) were identified, encoding basic phage-related functions, including phage DNA metabolic proteins, phage structural proteins, lysis-related proteins, and hypothetical proteins (**Fig. 5**). Genome annotations of KMM2 and KMM4 showed minor differences in their direct comparison, such as the length of genes and the presence or absence of specific genes (**Fig. 5B**), suggesting that these are two different and novel phages, verifying phage genome analysis using VIRIDIC.

**Figure 5.**
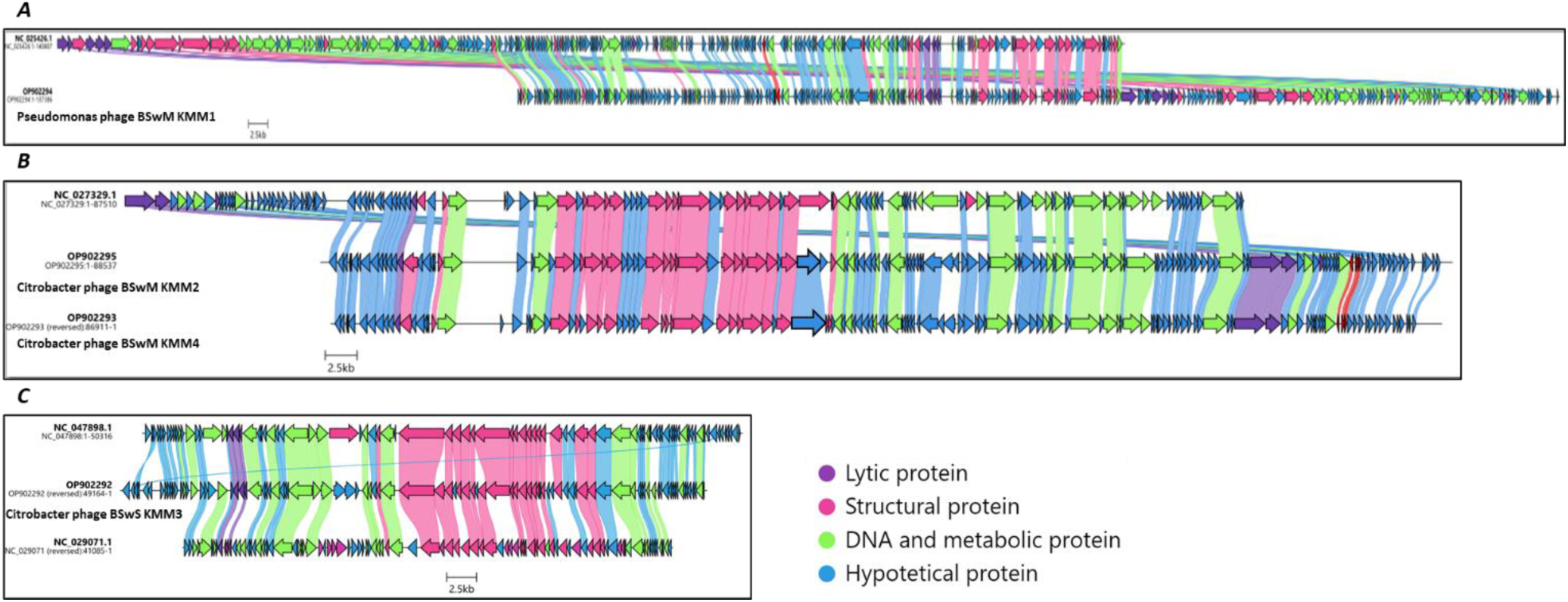
Genome annotations. The position and annotation of predicted viral genes in the phage genomes were visualized using Clinker. Coding domain sequences (CDS) are shown as arrows given the transcription direction; colors indicate their predicted function and amino acid sequence homologies to best homologs (nucleotide identity > 70 %) are presented by corresponding alignments. (***A***) KMM1 (OP902294), (***B***) KMM2 (OP902295) and KMM4 (OP902293), and (***C***) KMM3 (OP902292).

In summary, four new phages from the Baltic Sea water column were identified and characterized that efficiently and effectively infect *Pseudomonas* and *Citrobacter* bacteria, members of the complex microbiota of the moon jellyfish *A. aurita*. Particularly, phage KMM1 is a highly virulent, efficient phage infecting Gram-negative and Gram-positive bacterial species of genera *Pseudomonas* and *Staphylococcus*.

## Discussion

Viruses are found in all habitats on Earth (Abedon, 2008), but their importance is probably most evident in the ocean, where they are considered a source of diversity in genetic variation (Broniewski et al., 2020; Rousset et al., 2022; Weinbauer et al., 2003). Estimates suggest that phage numbers are tenfold higher than bacteria in the ocean, with phage particle estimates of 10^23^, resulting in turnover rates of 10^25^ infections and lysis events per second in the ocean impacting nutrient cycling (Breitbart et al., 2018; Jacquet et al., 2010). The relative proportions of lytic and lysogenic phages vary depending on various environmental factors, such as temperature, salinity, and nutrient availability (Munn, 2006; Zhang et al., 2022). In general, lytic phages tend to dominate in nutrient-rich environments with high microbial diversity and abundance and are thus more prevalent in surface waters, while lysogenic phages are more prevalent in nutrient-poor environments in deeper (oligotrophic) waters (Warwick-Dugdale et al., 2019). For instance, in the Baltic Sea, it has previously been demonstrated that lytic viruses are more common in surface waters, whereas lysogeny predominates in deep marine waters (Jakubowska-Deredas et al., 2012; Weinbauer et al., 2003). Lytic representatives of Siphovirus (52 %), Myovirus (42 %), and Podovirus (6 %) were consistent in the surface water throughout all seasons within the Baltic Sea (Jakubowska-Deredas et al., 2012; Nilsson et al., 2019; Von Scheibner et al., 2018; Šulčius & Holmfeldt, 2016). Those phages can have several roles particularly in the microbiomes of marine animals and plants, e.g., to maintain a healthy microbiome and prevent the spread of diseases (Brown et al., 2022; Mirzaei & Maurice, 2017; Pratama et al., 2020; Rolain et al., 2011). Further, phages appear responsible for several diseases that harm corals and their symbionts (Brüwer & Voolstra, 2018; Marhaver et al., 2008; Soffer et al., 2014; Work et al., 2021). Besides, phages promote the evolution of new traits or the acquisition of beneficial genes within a microbiome by mediating horizontal gene transfer between bacteria by transduction (Arnold et al., 2022; Juhas et al., 2009). Lastly, the impact mentioned above on nutrient cycling in marine ecosystems by breaking down bacterial cells and releasing nutrients into the environment can have important implications for marine organisms’ overall health and productivity (Middelboe, 2008). Although the role of Cnidarian bacterial communities has already been intensively investigated (Augustin & Bosch, 2010; Lee et al., 2018; Tinta et al., 2012; Weiland-Bräuer et al., 2020), the impact of (lytic) phages on Cnidarian and particularly on *A. auritás* bacterial colonizers has yet only rarely been studied. However, different *Hydra* species have been shown to harbor a diverse host-associated virome predominated by bacteriophages (Bosch et al., 2015; Deines et al., 2017; Grasis et al., 2014). Changes in environmental conditions altered the associated virome, increased viral diversity, and affected the metabolism of the metaorganism (Grasis et al., 2014). The specificity and dynamics of the virome point to a potential viral involvement in regulating microbial associations in the *Hydra* metaorganism (Bosch et al., 2015).

In this study, four phages (*Pseudomonas* phage KMM1, *Citrobacter* phages KMM2, KMM3, and KMM4) were isolated from the Baltic Sea water column (Kiel fjord) surrounding *A. aurita* individuals by a cultivation-based approach, infecting previously isolated bacteria, *Pseudomonas* and *Citrobacter*, both present in the associated microbiota of *A. aurita* (Weiland-Bräuer et al., 2015; Weiland-Bräuer et al., 2020). Both genera, *Pseudomonas* and *Citrobacter,* are Gram-negative bacteria widely distributed in marine environments, including seawater, sediments, and marine eukaryotes (Kimata et al., 2004; Lü et al., 2011; Ravi et al., 2018; Tkachenko et al., 2016). *Pseudomonas* species play a critical role in the marine nitrogen cycle, as they are capable of nitrogen fixation and denitrification (Zhang et al., 2020). While less is known about *Citrobacter*-specific ecological roles in marine environments, it can cycle degrading organic matter (Gatidou et al., 2022). Both *Pseudomonas* and *Citrobacter* are opportunistic pathogens and can cause infections in marine multicellular organisms (Lü et al., 2011; Thanigaivel et al., 2015; Tkachenko et al., 2016). However, they are likewise considered essential players in maintaining the health and balance of marine ecosystems (Marinho et al., 2009; Sun et al., 2020). Consequently, lytic phages infecting those bacteria might disturb the ecosystem and metaorganism homeostasis. Bacteriophages that infect *Pseudomonas* and *Citrobacter* have been identified in marine environments, and they may play essential roles in regulating the abundance and activity of these bacteria (Gatidou et al., 2022; Thanigaivel et al., 2015; Zhang et al., 2022). For example, bacteriophages that infect *Pseudomonas* can control its population size and limit its ability to degrade organic matter, significantly impacting nutrient cycling in marine ecosystems (Knezevic et al., 2011; Wommack & Colwell, 2000). Similarly, bacteriophages that infect *Citrobacter* can reduce the abundancy of this bacterium and thus limit its ability to colonize and persist in marine environments, potentially reducing the risk of animal diseases and human exposure to this pathogen, e.g., through seafood consumption (Dy et al., 2018; Edeh & Nsofor, 2023; Tkachenko et al., 2016).

KMM1 reflects a broad host range phage capable of infecting a wide range of bacterial hosts, whereas KMM2/3/4 reflect narrow host range phages, infecting only a limited number of bacterial strains or species (Breitbart et al., 2018; Ross et al., 2016). Remarkably, KMM1 efficiently infects Gram-negative *Pseudomonas* strains as well as a variety of strains of various Gram-positive *Staphylococcus* species. Gram-negative and Gram-positive bacteria differ in cell walls and membrane characteristics (Dziarski & Gupta, 2000; Elson et al., 2007). Gram-positive bacteria have thick multilayered peptidoglycan in their cell envelope, while Gram-negative bacteria’s cell walls have just a thin layer of peptidoglycan covered by an outer membrane. Consequently, phages require strain-, species-, or even higher order– specific characteristics for bacteria recognition, attachment, and lysis. Phages usually use their tail fibers or spikes to recognize and attach to specific receptors on the bacterial cell wall of Gram-negatives, such as lipopolysaccharides (LPS) or outer membrane proteins (Datta et al., 1977; Proft & Baker, 2009). The tail fibers or spikes bind to these receptors through specific interactions, which can be highly selective. Gram-positive bacteria are likewise precisely recognized and infected by teichoic acids or other surface proteins (de Carvalho, 2016; Frias, 2011; Mahony et al., 2016). Moreover, phages replicate differently inside their host cells, depending on the bacterial structures. Gram-negative-specific phages enter the host cell by injecting their genetic material directly into the cytoplasm, where it replicates and assembles new phage particles (Zampara et al., 2020). In contrast, Gram-positive phages usually enter the host cell by binding to receptors on the host surface, replicating and assembling new phage particles in the cytoplasm (Dunne et al., 2018; Perea et al., 2021). To the best of our knowledge, few characterized phages in the marine environment have such a broad host range as the phage KMM1. However, four characterized phages are already isolated from activated sludge samples, and one phage identified from a freshwater sample capable of infecting both genera (Khan et al., 2002; Lin et al., 2011; Wang et al., 2020). We can only speculate that KMM1 might recognize and bind to conserved structural components of bacterial cells. KMM1 might has evolved in the marine environment due to the frequent presence of *Pseudomonas* and *Staphylococcus* allowing recognition and infection of a wide range of bacterial hosts. Potentially, KMM1 prefers the highly abundant *Pseudomonas* as a host, but it can also infect *Staphylococcus* under certain, possibly stressful, environmental conditions that promote *Staphylococcus* abundance. Genome analysis of KMM1 supports the experimentally collected data, i.e., that it is capable of infecting both Gram-positive and Gram-negative bacteria. We identified several features within the KMM1 genome indicative of Gram- negative bacteria infection, such as *Pseudomonas*. In more detail, we identified YbiA-like superfamily proteins (IPR037238) derived from various Gram-negative bacteria, such as *E. coli* K-12 (Quin et al., 2012), and *Pseudomonas,* suggesting the infection of Gram-negative bacteria. Similarly, the invasin/intimin cell-adhesion domain (IPR008964) was found in the phage genome, common in phages that infect Gram-negative bacteria such as *Erwinia* (Born et al., 2011). In contrast, we further identified the CHAP (cysteine, histidine-dependent amidohydrolase/peptidase) domain (IPR007921) in the KMM1 genome. This domain has been shown to be specifically responsible for the major catalytic activity of the endolysin in degrading cell wall peptidoglycan of staphylococci, including methicillin- resistant *Staphylococcus aureus* (Fenton et al., 2011; O’flaherty et al., 2005; Sanz-Gaitero et al., 2013). Future studies will reveal more insights into the complex attachment and infection mechanisms of phage KMM1 and will disclose the underlying mechanisms of its broad host range.

All phages identified in this study showed effective and efficient lysis of *Pseudomonas* and *Citrobacter* by short latency periods and high burst sizes (**Table 3**). Those characteristics are important features affecting natural microbiomes and relevant for potential therapeutic applications. Phage therapy uses intact natural phages or phage compounds to treat bacterial infections (Nilsson, 2014). Due to the growing number of antibiotic-resistant bacterial species and the ban on the use of antibiotics in the aquatic environment (Broniewski et al., 2020; Chuah et al., 2016; Frieri et al., 2017; Hossain et al., 2022), the interest in phage therapy particularly for aquaculture increased during the last few decades (Culot et al., 2019; Nakai & Park, 2002; Ramos-Vivas et al., 2021). Phage therapy relies on phages’ extraordinary qualities, including host specificity, self-replication, wide distribution, and safety (Hyman, 2019; Nilsson, 2014; Ramos-Vivas et al., 2021; Skurnik et al., 2007). Since phages are a natural way of managing bacterial infections, their usage does not contribute to the development of antibiotic resistance or the deposition of harmful residues in the environment. Finally, phages are versatile since they may be used alone or in cooperation with antibiotics or other therapies to improve their potency against bacterial infections. These features are entirely applicable in aquaculture, where traditional approaches to deal with pathogenic bacteria, such as antibiotics, are impossible (Dy et al., 2018; Edeh & Nsofor, 2023). Particularly the identified broad host range bacteriophage KMM1 targets various bacteria, increasing its usefulness in aquaculture where different bacterial species can cause infections (Hyman, 2019). KMM1’s high infectivity rate can quickly and effectively reduce bacterial populations, limiting the spread of infection. This phage’s stability, safety, and cost-effective production under different environmental conditions, such as pH and temperature, must be tested for potential use as an effective alternative to antibiotics in controlling bacterial infections in aquaculture.

Lastly, our study demonstrated that phage research methods are still in their infancy, although several benchmarking studies on genomic and seasonal variation and diversity of tailed phages in the Baltic Sea were already published (Jakubowska-Deredas et al., 2012; Nilsson et al., 2019). Bioinformatics tools specifically developed explicitly for phages still lag behind similar tools used in bacterial research (Coclet & Roux, 2021; Martinez-Vaz & Mickelson, 2020; Schackart III et al., 2023). However, some improved tools in recent years include Phaster, a web-based tool for identifying and annotating phage sequences in bacterial genomes and predicting their completeness (Arndt et al., 2016). VirSorter, VirFinder, and Phage AI implement machine learning to identify viral sequences in metagenomic datasets, distinguish between viral genomes, plasmids, and transposons, and predict phage host ranges and other characteristics (Fang et al., 2019; Lenneman et al., 2021). In the present study, we used VICTOR for pairwise distance-based comparisons of the whole genome, vConTACT2 to determine the genus-level taxonomy of viral genomes, and VIRIDIC for calculating the intergenomic distance of viruses. The results obtained on species assignment of KMM2 and KMM4 using VICTOR and vConTACT2 demonstrate that phage softwares result in contrasting statements and must be improved. Using VICTOR and vConTACT2 did not allow for the differentiation of highly similar phages. Experimental data on differences within the lytic cycle and host range (**Fig. 2 and 3**), in combination with genome annotation, pointed to different species assignments of KMM2 and KMM4 (**Fig. 5**). However, by improving estimates of phage genome similarity, particularly for distantly related phages, analyzing datasets including thousands of phage genomes, and creating an informative heatmap that includes not only the similarity values but also information about the genome lengths and aligned genome fraction, VIRIDIC finally confirmed species assignments.

In summary, the present study describes the identification and characterization of four novel bacteriophages. The broad-host-range phage KMM1 infects members of the genera *Pseudomonas* and *Staphylococcus*. Likely, KMM1 adapted its attachment and infection mechanisms through co-evolution with its bacterial hosts. The phages KMM2-4 infect *Citrobacter* and close relatives of *Enterobacteriaceae,* thus possessing a narrow-host range. Although all identified phages demonstrated effective and efficient lytic properties relevant for phage application, future studies on the impact of those phages on the native microbiome of the moon jellyfish *A. aurita,* a model in metaorganism research, are of particular interest. Such studies may provide insights into the complex interdependence of phages and their bacterial hosts and how these relationships affect microbiomes to investigate the impact on the eukaryotic host *A. aurita*. It is conceivable that the infection of *A. aurita* with the phages KMM1-4 might cause substantial changes in the bacterial community, potentially disrupting the multicellular host’s homeostasis. Even assuming that the phages impact *A. auritás* microbiome structure, it may be that the eukaryotic host has mechanisms to maintain its homeostasis. Future studies will focus on the characterization of attachment and infection mechanisms, particularly of phage KMM1, and the impact of the identified phages on the microbiome and, consequently, the health of *A. aurita*.

## Author Contributions

Conceptualization, R. A. S. and N. W.-B.; methodology, M. S.; investigation, M. S.; formal analysis, M. S.; microscopy and micrographs analysis, U. R. and M. B. ; bioinformatics analysis, C. M. C. and A. W.; writing—original draft preparation, M. S., N. W.-B., and R. A. S.; writing— review and editing, all authors; supervision, N. W.-B. and R. A. S.; project administration, R. A. S.; funding acquisition, R. A. S.. All authors have read and agreed to the published version of the manuscript.

## Acknowledgments

We thank the Institute of Clinical Molecular Biology in Kiel, Germany, for providing Sanger sequencing. We acknowledge financial support by Land Schleswig-Holstein within the funding program Open Access Publikationsfonds.

## Data Availability Statement

The phage genomes are available at NCBI under the accession numbers OP902294: Pseudomonas phage BSwM KMM1, OP902295: Citrobacter phage BSwM KMM2, OP902292: Citrobacter phage BSwS KMM3, and OP902293: Citrobacter phage BSwM KMM4. Sequences were deposited on the Sequence Read Archive (SRA) under BioProject PRJNA908753 and raw sequences accession numbers SRR22580853, SRR22580850, SRR22580849, and SRR22580845. Bacterial sequence data derived from Sanger sequencing of the 16S rRNA genes of bacterial isolates is deposited under GenBank accession numbers OQ397638-OQ397653, OQ398153-OQ398172.

